# De novo administration of antiviral monoclonal antibodies against SARS-CoV-2 or influenza using mRNA lipid nanoparticles

**DOI:** 10.1101/2025.07.23.666463

**Authors:** Mai N. Vu, Jessica A. Neil, Charley Mackenzie-Kludas, Andrew Kelly, Hyon Xhi Tan, Kanta Subbarao, Wen Shi Lee, Adam K. Wheatley

## Abstract

Monoclonal antibodies (mAbs) are an emerging class of therapeutics for the prevention and treatment of viral infections. Recent advances in mRNA/lipid nanoparticle (LNP) technology provide a potential new modality for the expression of mAbs in vivo, potentially bypassing the need for recombinant manufacturing of mAb proteins. In this study, we compared traditional infusion of neutralising mAbs targeting SARS-CoV-2 or influenza to mRNA-based induction of de novo mAb expression in treated mice. High serum concentrations of mAbs were achieved upon delivery of a single mRNA encoding both heavy and light chains via intravenous or intramuscular routes using prototypic LNP formulations. However, pharmacokinetics were heavily influenced by the induction of anti-drug antibody responses directed against the encoded mAbs, driving reductions in in vivo half-life and compromising protective capacity against SARS-CoV-2 Omicron BA.1 infection. Overall, mRNA/LNP delivery comprises a feasible and attractive pathway to speed the development and deployment of antiviral antibodies, however optimisation of LNP formulation, dosing and administration routes is required to maximise protective potential.

## INTRODUCTION

Viral pathogens constitute a persistent threat to global health and economic security. Neutralising antibodies, which can directly block infection, are a key correlate of immune protection^1,2^ and highly potent neutralising mAbs have been developed as effective antiviral drugs for Ebola^3^, RSV^4^ and SARS-CoV-2^5^. However, for endemic viruses, the high production costs historically associated with mAb-based treatments have been a barrier to development, while in the context of outbreaks or pandemics, the long lead times required for cell-based recombinant manufacture significantly delay the potential introduction of mAbs to protect vulnerable populations.

Recent technological advances in the delivery of mRNA encapsulated within lipid nanoparticles (LNP) have transformed the delivery of vaccines, with products from Pfizer/BioNtech^6^ and Moderna^7^ demonstrating considerable clinical impact during the recent COVID-19 pandemic. However, the potential for mRNA/LNPs in the delivery of antibody-based drugs is relatively understudied, with optimal parameters for mRNA design, LNP formulation, and route of administration yet to be comprehensively defined. Intravenous (i.v.) administration of LNPs co-encapsulating two separate mRNAs encoding the heavy (HC) and light immunoglobulin chains (LC) has achieved marked serum concentration of mAbs against SARS-CoV-2^8^ or SFTSV^9^ in pre-clinical models. In a clinical study, mRNA/LNP delivery of a mAb targeting Chikungunya virus was safe in humans and similarly induced robust serum titres.^10^ However, optimal LNP formulations, mRNA formats, and administration routes for efficient delivery of mRNA-encoded mAbs to both systemic and mucosal compartments require further exploration, in particular for respiratory viruses where mucosal localisation may be crucial to prevent viral entry, replication, and transmission.^11,12^

Here, we contrasted mRNA/LNP versus conventional delivery of two potent neutralising human IgG mAbs against COVID-19 (PDI204) and influenza (HV-B10).^13,14^ A single mRNA construct encoding both HC and LC drove robust mAb expression in mice and readily detectable serum neutralising titres. Classical LNP formulations analogous to the Pfizer/BioNTech COVID-19 vaccine (Comirnaty) elicited high mAb titres within the blood and lung after i.v. or intramuscular (i.m.) administration, but not following intranasal (i.n.) instillation. In comparison to recombinant protein controls, mRNA-delivered mAbs elicited comparable peak serum concentrations, but were subject to more rapid waning, likely attributable to the induction of anti-drug antibodies (ADA) heightened by the adjuvant properties of LNPs. Finally, in experimental challenge models of influenza or SARS-CoV-2, we found mRNA/LNP delivery of mAbs could provide a degree of protection against the development of severe disease, although protective efficacy was modulated by dose, pathogen, and timing of challenge. Our study highlights mRNA/LNPs constitute a tractable pathway for the rapid and efficient delivery of protective antibodies against viral diseases.

## RESULTS

### Expression of functional monoclonal antibody in vitro and in vivo following mRNA/LNP delivery

Previously, we isolated a potent SARS-CoV-2 neutralising human IgG mAb PDI204, which targets the receptor binding domain (RBD) of the spike protein.^13^ The safety and efficacy of this mAb in humans are being evaluated in a phase I clinical trial (NCT06965751). We explored the potential of mRNA/LNPs to deliver PDI204 using two mRNA formats: (i) encoding the HC and LC separately in two mRNA constructs, or (ii) combining the two chains in a single mRNA construct linked by a furin cleavage site and self-cleaving peptide P2A motif^15^ (HC-P2A-LC). Using a formulation analogous to Comirnaty (ALC-0315) (Table S1),^16^ we prepared LNPs carrying an admix of HC and LC mRNAs (at 1:1 molar ratio), or HC-P2A-LC mRNAs for in vitro and in vivo delivery (**Figure 1a**). Similar physicochemical properties (diameters of ∼ 60 nm, negative charges of approximately – 8 mV, and encapsulation efficiencies of ∼ 90%) were observed between these two LNPs (Table S2).

**Figure 1:**
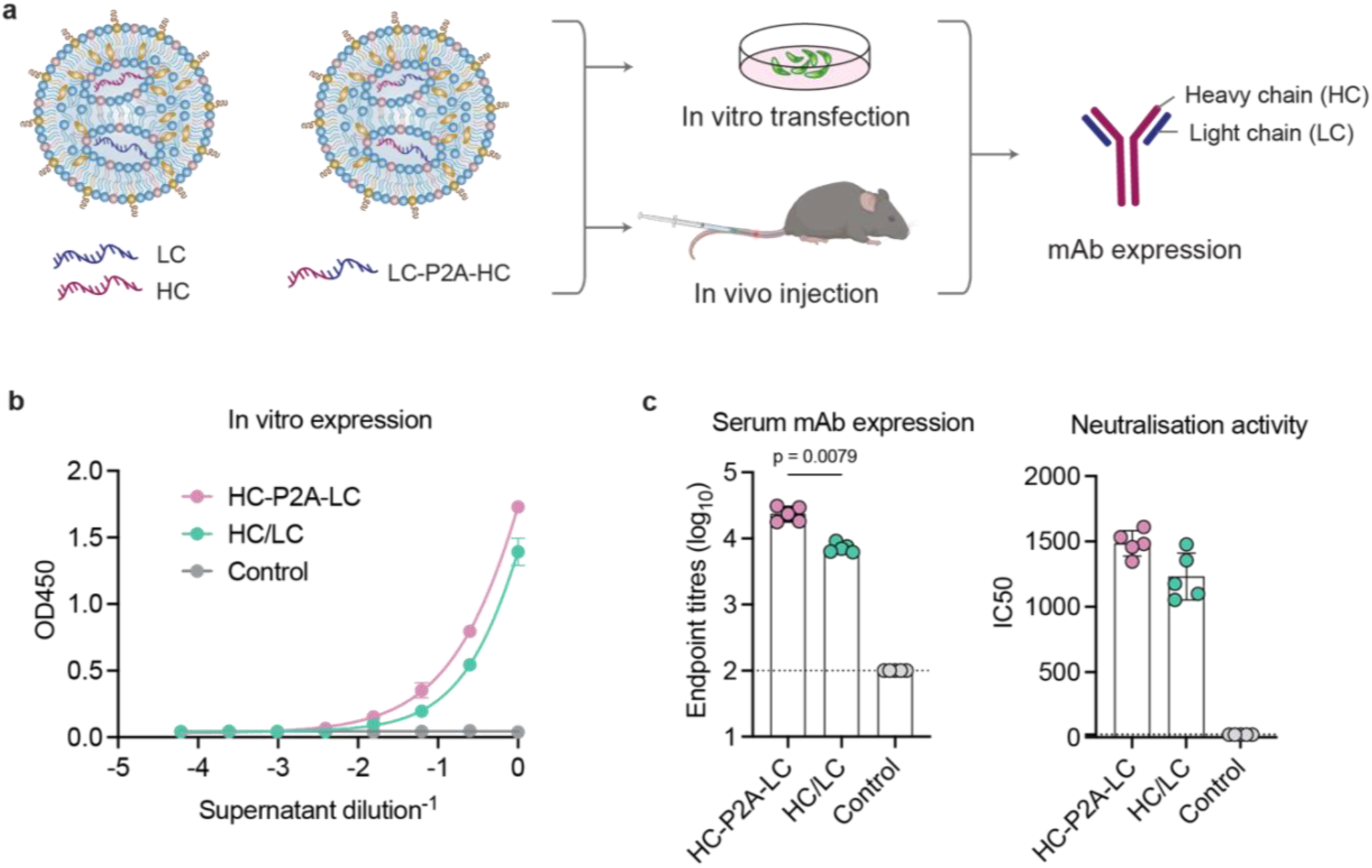
Rational design and characterisation of mRNA/LNPs delivering monoclonal antibodies. **a**). LNPs delivering PDI204 HC and LC encoded in two separate mRNAs or in a single mRNA with a P2A linker were transfected in HEK293T cells or i.v. injected into C57BL/6 mice (n = 5 per group). LNPs carrying ovalbumin mRNA were used as a negative control. **b**). PDI204 expression in HEK293T cells at 48h after transfection using an ELISA assay against the SARS-CoV-2 spike protein. **c**). Titres of PDI204 expression in mouse serum at 48h post-injection measured by ELISA assay against SARS-CoV-2 spike protein (left) and neutralisation activities of the mouse sera against ancestral SARS-CoV-2 viruses using a microneutralisation assay (right). Data are shown as median ± IQR. Statistical significance was determined by a Kruskal-Wallis test followed by post-hoc Dunn’s multiple comparisons test.

The expression of functional PDI204 mAbs following mRNA/LNP transfection of HEK293T cells was confirmed by ELISA, with no marked differences in the concentrations of mAbs expressed in supernatants between the two mRNA formats (Figure 1b). However, at 48 hours after i.v. administration into C57BL/6 mice, the combined HC-P2A-LC mRNA/LNPs produced significantly higher serum PDI204 titres (*p* = 0.0079), and greater neutralisation activity compared to delivery of the admixed HC/LC mRNAs (Figure 1c). Overall, both delivery approaches demonstrated the capacity to efficiently deliver functional PDI204 mAbs in cell culture and mice, with the combined HC-P2A-LC mRNA construct selected for the additional studies.

### Effects of LNP formulation and administration routes on the biodistribution of mRNA-delivered mAbs

Positively charged LNPs formulations have been reported to enhance mRNA delivery to the lungs following i.v. injection,^17,18^ however our previous studies have highlighted both anionic and cationic LNPs can efficiently deliver mRNA to the respiratory mucosa following i.n. administration.^16^ To assess the effects of LNP formulation and delivery routes on mAb biodistribution, we utilised these two LNP formulations developed previously^16^ to encapsulate PDI204 HC-P2A-LC mRNA: (i) an anionic, Comirnaty-like ALC-0315 LNP, and (ii) a cationic MC3-DOTAP LNP (Table S1). Compared to ALC-0315 (∼ 64.1 nm, –8.09 mV), the MC3-DOTAP LNPs had a larger diameter of ∼121.7 nm and a positive charge of +15.2 mV.

PDI204 ALC-0315 and MC3-DOTAP mRNA/LNPs were delivered into mice via i.v., i.n., and i.m. injection. PDI204 expression in mouse serum, bronchoalveolar-lavage fluid (BALF), and nasal washes was quantified at 48h post administration (**Figure 2a**). I.v. and i.m. delivery of ALC-0315 LNPs drove the accumulation of high levels of PDI204 in serum, BALF, and nasal wash samples compared to animals treated with control mRNA/LNPs (Figure 2b), but minimal expression was detected at any site following i.n. delivery. In contrast, MC3-DOTAP LNPs were poor at driving mAb expression in vivo, although there was evidence of some expression within the lungs following i.n. administration. Overall, ALC-0315-based formulations and i.v. and i.m. routes appeared best suited to deliver mAbs in vivo.

**Figure 2:**
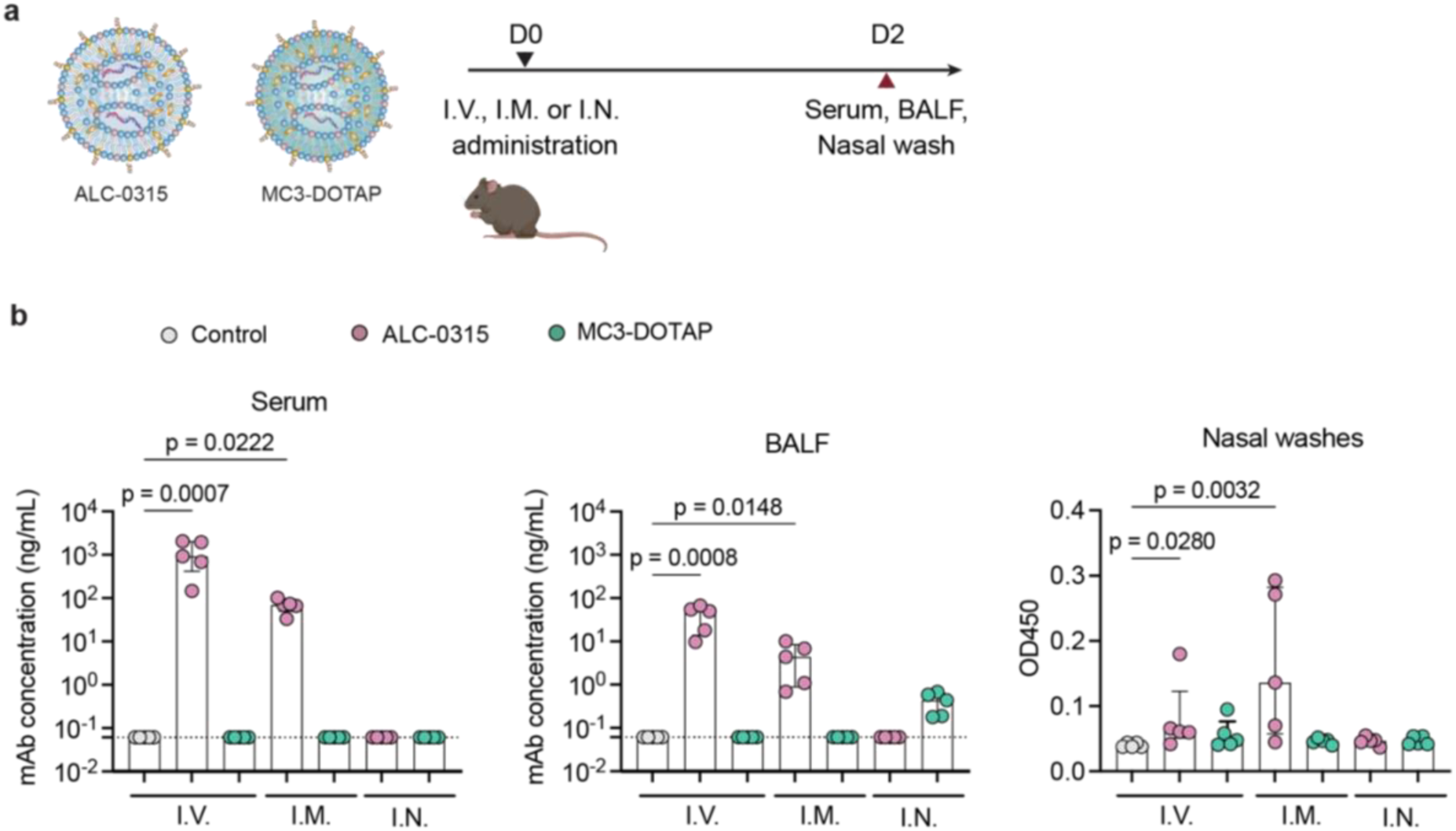
Effects of LNP formulations and administration routes on biodistribution of mRNA-delivered mAbs in vivo. **a**). ALC-0315 and MC3-DOTAP LNPs delivering PDI204 HC-P2A-LC mRNAs at 5 μg were injected into C57BL/6 mice (n = 5) via either i.v., i.m., or i.n. routes. At 48h post-injection, serum, BALF, and nasal wash samples from the injected mice were collected. **b**). Levels of PDI204 in serum and mucosal washes were measured by ELISA against SARS-CoV-2 spike protein. Data are shown as median ± IQR. Statistical significance was determined by a Kruskal-Wallis test followed by post-hoc Dunn’s multiple comparisons test.

### Pharmacokinetics and protective capacity of mRNA-delivered mAbs against SARS-CoV-2

We next compared the serum pharmacokinetics (PK) of mAbs delivered via ALC-0315 mRNA/LNPs versus recombinant proteins after i.v. administration (**Figure 3a**). Both mRNA and recombinant protein were administered at 10, 5, and 1 μg doses per animal and monitored over a 70-day sampling period. Peak serum concentrations of PDI204 were detected at day 1 after administration of recombinant protein, while for mRNA/LNPs we generally observed serum concentrations continued to increase out to days 3-5 (Figure 3b). Both protein and mRNA displayed similar dose-response relationships, however we observed marked differences in their persistence. While recombinant PDI204 proteins remained detectable up to day 70 when given at 5 or 10 μg doses, serum concentrations of mRNA-delivered mAbs rapidly declined from 7 days post-administration and were generally cleared by day 28 (Figure 3b). PK dynamics are summarised in **Table 1**.

**Figure 3:**
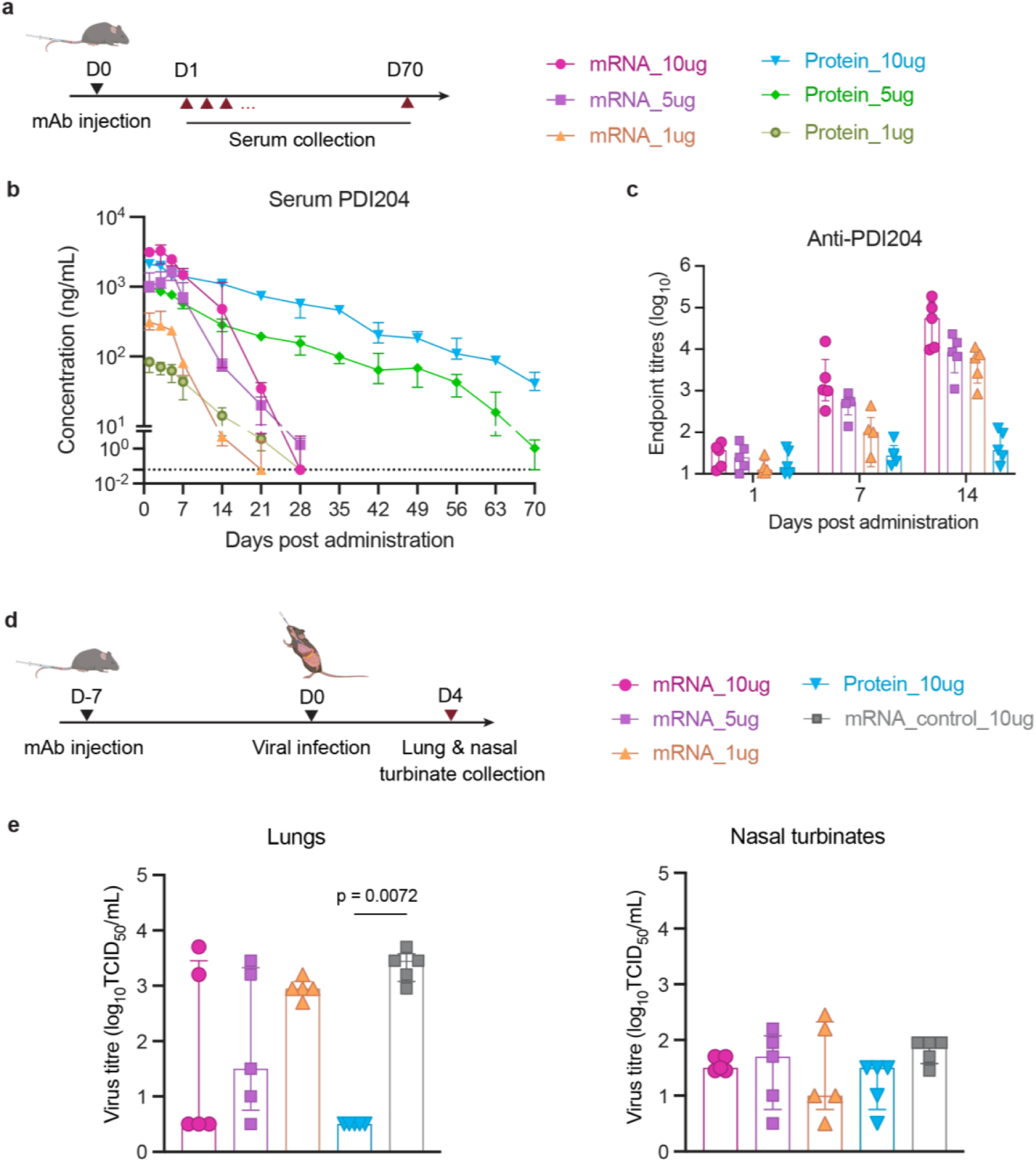
PK profile and prophylactic efficacy of PDI204 mRNA/LNPs against SARS-CoV-2 infection. **a**). PDI204 mRNA/LNPs or proteins at 10, 5, or 1 μg were i.v. injected into C57BL/6 mice (n = 5). Sera from injected mice were collected at 1, 3, 5, 7, 14, 21, 28, 35, 42, 49, 56, 63, and 70 days post-injection. **b**). PDI204 concentrations in the sera over time were calculated using ELISA against SARS-CoV-2 spike protein. **c**). Anti-PDI204 antibody titres in mouse serum at days 1, 7, and 14 post-injection were measured by ELISA assay. **d**). k18-hACE2 mice (n = 5) were i.v. injected with either PDI204 mRNA/LNPs at 10, 5, or 1 μg or proteins at 10 μg. Irrelevant mAb mRNA/LNPs (10 μg) were used as a negative control. At day 7 post mAb injection, the mice were i.n. infected with SARS-CoV-2 BA.1 variants. **e**). Viral titres within lung and nasal turbinate homogenates were quantified by a TCID50 assay at day 4 post-infection. Data are shown as median ± IQR. Statistical significance was determined by a Kruskal-Wallis test followed by post-hoc Dunn’s multiple comparisons test.

**Table 1:** PK parameters for serum PDI204 delivered by mRNA/LNPs and protein.

| PDI204 | Half-life (day) |  | T<sub>max</sub><br>(day) | C<sub>max</sub><br>(µg/mL) | AUC<sub>last</sub><br>(day*µg/mL) |
| --- | --- | --- | --- | --- | --- |
|  | Before day 7 | After day 7 |  |  |  |
| mRNA_10µg | 7.7 ± 1.2 | 1.8 ± 0.5 | 2.5 ± 1.0 | 3.4 ± 0.6 | 26.6 ± 7.2 |
| mRNA_5µg | 8.1 ± 3.1 | 2.3 ± 0.6 | 5.0 | 1.7 ± 0.5 | 11.6 ± 3.5 |
| mRNA_1µg | 3.2 ± 0.8 | 2.0 ± 0.9 | 2.6 ± 0.9 | 0.3 ± 0.1 | 1.9 ± 0.3 |
| Protein_10µg | 10.4 ± 4.0 | 12.7 ± 2.5 | 1.8 ± 1.1 | 2.2 ± 0.3 | 39.2 ± 5.1 |
| Protein_5µg | 9.7 ± 2.9 | 11.5 ± 2.7 | 1.0 | 1.0 ± 0.07 | 13.2 ± 0.9 |
| Protein_1µg | 6.4 ± 2.3 | 3.0 ± 1.2 | 1.0 | 0.08 ± 0.02 | 0.6 ± 0.2 |

Comirnaty formulation was developed for vaccine delivery, which has been demonstrated to induce strong immune responses to the encoded antigen^6^, with the ionisable lipid ALC-0315 comprising a robust immune adjuvant.^19^ We therefore assessed the induction of anti-drug antibodies (ADA; anti-PDI204) in mouse serum at days 1, 7 and 14 post administration (Figure 3c). In contrast to mice administered a high dose of PDI204 protein (10 μg), all animals given mRNA/LNPs developed a significant antibody response against PDI204 over time. The findings suggest the rapid reduction in serological titres of mRNA-delivered PDI204 seen from day 7 post-administration may be attributed to ADA responses, with ADA induction likely precipitated by the immunogenic LNP delivery system.

We next assessed the prophylactic efficacy of PDI204 delivered by mRNA/LNPs in a transgenic human ACE2 mouse model (k18-hACE2).^13^ At day 7 post i.v. injection of mAbs, the mice were intranasally infected with SARS-CoV-2 Omicron BA.1 variant (strain SARS-CoV-2/Australia/NSW/RPAH-1933/2021) (Figure 3d). As the BA.1 strain causes limited pathology or weight loss in this model, we measured viral titres in the lungs and nasal turbinates at day 4 post-infection, corresponding to day 11 post-mAb administration (Figure 3e). Treatment with 10 μg PDI204 protein resulted in robust protection of the lungs, with no viral loads detected post challenge (*p* = 0.0072). For animals receiving mRNA/LNPs, we observed more animal-to-animal variability, but there was a dose-proportional reduction in viral loads associated with increasing treatment doses from 1 to 10 μg. Small and comparable amounts of virus were detected within the nasal turbinates of all challenged animals, likely reflective of the initial inoculum. Overall, the challenge study indicates that mRNA/LNP delivery of anti-viral mAbs can afford a degree of protection against SARS-CoV-2, although this appears inferior in this model compared to administration of recombinant protein, which was likely impacted by the development of ADA over the extended 11-day challenge window.

### Influenza mAbs delivered by mRNA/LNPs protect mice against lethal viral challenge

Influenza is a highly infectious respiratory pathogen that causes seasonal epidemics and periodic pandemics. We therefore explored the potential of mRNA/LNP delivery of a human anti-influenza mAb and assessed its protection against lethal influenza infection. HV-B10 was selected as a prototypic potent neutralising mAb targeting the hemagglutinin (HA) on influenza pH1N1 viruses^14^. The HV-B10 mAb mRNA was also designed with heavy and light chains linked by a P2A motif.

The PK profile of HV-B10 delivered using ALC-0315 mRNA/LNPs was compared to recombinant protein as before (**Figure 4a**). Similar to PDI204, we found administration of HV-B10 protein drove a rapid rise in serum concentration of mAb, followed by an extended decay period out to more than 50 days for animals receiving 5 or 10 μg doses (Figure 4b). In contrast, mRNA-delivered HV-B10 rapidly achieved high serum concentrations, but were subject to rapid clearance from day 7 onwards (**Table 2**). This again likely corresponded with the induction of anti-HV-B10 antibodies, which were readily evident in mice at days 7 and 14 post mRNA/LNP administration (Figure 4c). Serum concentrations of HV-B10 after mRNA delivery peaked at higher levels than corresponding doses of recombinant protein and at levels higher than seen with PDI204, suggesting that the specific characteristics of each mRNA construct can influence overall delivery efficiency in vivo.

**Figure 4:**
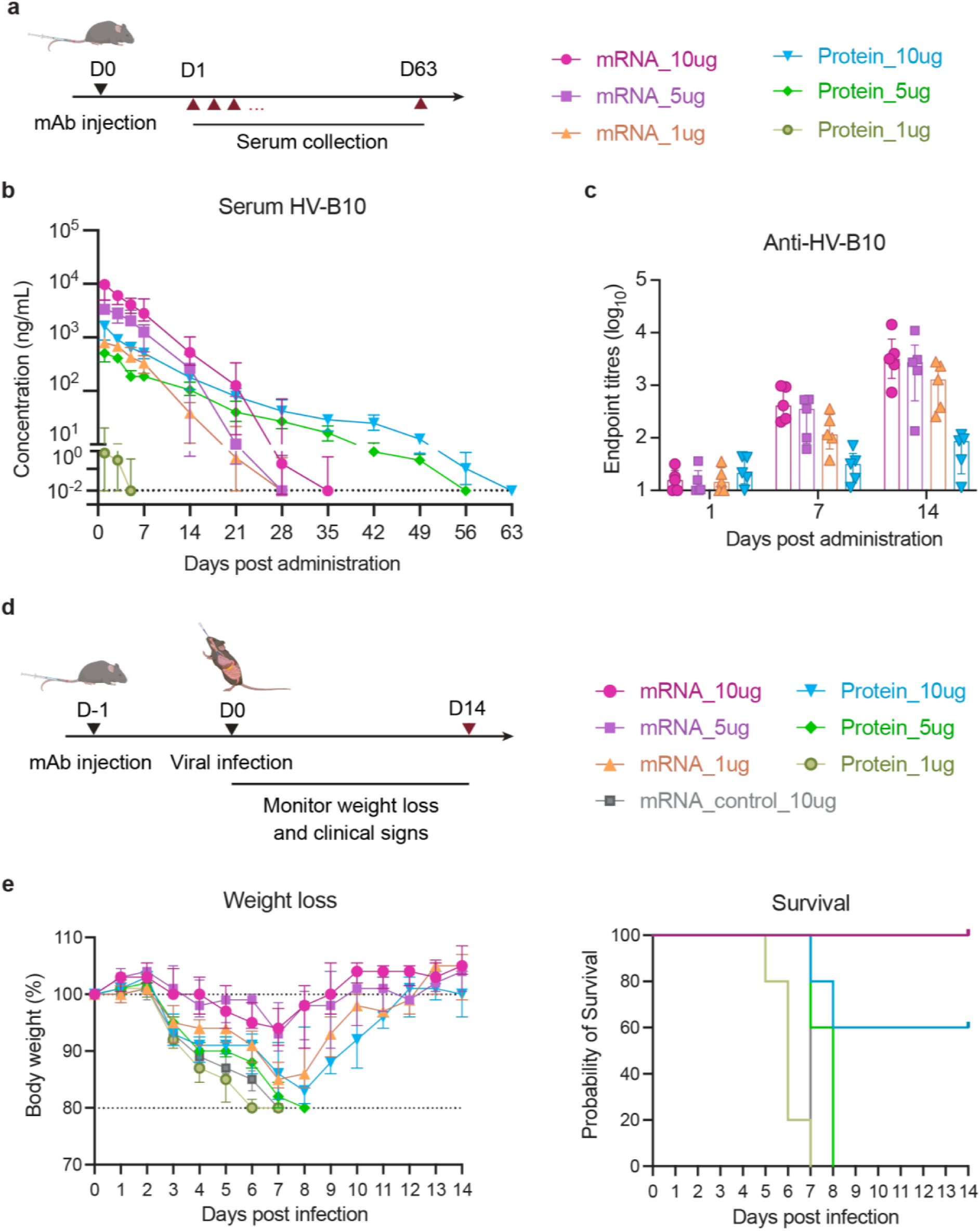
PK profile and protection efficacy of HV-B10 mRNA/LNPs against influenza infection. **a**). HV-B10 mRNA/LNPs or proteins at 10, 5, or 1 μg were i.v. injected into C57BL/6 mice (n = 5). Sera from injected mice were collected at 1, 3, 5, 7, 14, 21, 28, 35, 42, 49, 56, and 63 days post injection. **b**). HV-B10 concentrations in the sera over time were calculated using ELISA against A/California/04/2009 HA protein. **c**). Anti-HV-B10 antibody titres in mouse serum at days 1, 7, and 14 post-injection were measured by an ELISA assay. **d**). At 24h post mAb injection, the mice were i.n. infected with 100 pfu of A/California/04/2009 virus. Irrelevant mAb mRNA/LNPs (10 μg) were used as a negative control. **e**). Changes in body weight (left) and **s**urvival rates (right) of injected mice in 14 days post-viral challenge. Data are shown as median ± IQR.

**Table 2:** PK parameters for serum HV-B10 delivered by mRNA/LNPs and protein.

| HV-B10 | Half-life (day) |  | T<sub>max</sub><br>(day) | C<sub>max</sub><br>(µg/mL) | AUC<sub>last</sub><br>(day*µg/mL) |
| --- | --- | --- | --- | --- | --- |
|  | Before day 7 | After day 7 |  |  |  |
| mRNA_10µg | 5.3 ± 2.4 | 2.4 ± 1.1 | 1.4 ± 0.9 | 8.7 ± 2.9 | 49.5 ± 19.1 |
| mRNA_5µg | 3.9 ± 1.5 | 1.8 ± 0.6 | 1.0 | 3.8 ± 1.2 | 21.3 ± 7.6 |
| mRNA_1µg | 4.6 ± 1.5 | 2.4 ± 1.0 | 1.8 ± 1.1 | 0.8 ± 0.1 | 5.0 ± 1.0 |
| Protein_10µg | 6.7 ± 3.2 | 6.5 ± 0.7 | 1.4 ± 0.9 | 1.5 ± 0.5 | 9.4 ± 0.9 |
| Protein_5µg | 5.3 ± 1.2 | 7.2 ± 0.8 | 1.4 ± 0.9 | 0.5 ± 0.1 | 4.0 ± 0.8 |
| Protein_1µg | 1.3 ± 0.6 | - | 1.0 | 0.02 ± 0.01 | 0.03 ± 0.02 |

The protective capacity of HV-B10 delivered as mRNA/LNPs or proteins was assessed in animals intranasally challenged with a lethal dose (100 pfu) of A/California/04/2009 virus (Figure 4d).^20^ Mice receiving HV-B10 as recombinant protein showed a degree of protection at a 10 μg dose (approximately equivalent to 0.5 mg/kg in a ∼ 20-g animal) with delayed weight loss and 60% survival (Figure 4e). Doses of 5 μg and 1 μg of HV-B10 protein were not protective relative to controls. In contrast, all mice receiving HV-B10 mRNA at 10, 5, or 1 μg doses were protected from viral challenge, with a dose-associated reduction in weight loss observed. Overall, HV-B10 was efficiently delivered by mRNA-LNPs, which afforded effective protection against a lethal influenza virus challenge.

## DISCUSSION

While vaccination represents a key tool for population-level protection from viral infection, antiviral therapeutics such as mAbs are important for supplemental protection of vulnerable subpopulations (e.g. immunocompromised or elderly individuals). mRNA/LNP technologies demonstrated a rapid design, development and deployment capability during the recent COVID-19 pandemic that could be applied to mAb-based antivirals. Here, we tested mRNA/LNP delivery platforms for two antiviral mAbs in pre-clinical models of infection. In line with previous reports in pre-clinical^8,9^ and clinical studies^10^, we found co-formulation of HC and LC mRNA cargoes could efficiently drive mAb expression in vitro and in vivo, although the use of a single mRNA with HC and LC linked by P2A peptides was equally effective and potentially simpler to manufacture and formulate.

The lipid components selected for LNP formulation have shown marked impacts on the efficiency of mRNA delivery, trafficking and immunostimulatory activity.^21^ A limited number of LNP formulations have advanced to human clinical use, although compositions with their ionisable lipids such as ALC-0315 (BioNTech/Pfizer), SM-102 (Moderna) and Dlin-MC3-DMA (MC3; Onpattro/Alnylam Pharmaceuticals) are most extensively studied. We find Comirnaty-like formulations could drive efficient mAb accumulation in vivo after i.v. and i.m, but not i.n. administration, while MC3 formulations with DOTAP included, previously reported to enhance delivery to lung tissues^17,18^, were demonstrably inferior and failed to robustly induce mAb expression in treated mice. The ionizable lipid component (ALC-0315) used in vaccine platforms such as the Comirnaty COVID-19 vaccine is known to possess strong adjuvant properties.^19^ As a consequence, we found this formulation drove the rapid elicitation of ADA against the antiviral human mAbs expressed in this study. This in turn dramatically compromised the in vivo half-life, with rapid clearance observed of serum mRNA-delivered PDI204 or HV-B10 relative to recombinant protein controls. ADA and accelerated clearance are likely to significantly affect the protective capacity of these mAbs during pathogenic challenge. In the context of SARS-CoV-2, when animals were challenged relatively late (11 days) after mRNA/LNP administration, protection was inferior to animals given PDI204 recombinant protein, which displayed sterilising immunity in the lungs. In contrast, when animals were challenged with influenza but only 1 day post mRNA/LNP administration, protection from mRNA-expressed HV-B10 was high and potentially more robust than that observed with recombinant protein controls. Although immune responses to mRNA-encoded mAbs will be heavily diminished when antibodies from matched species are used, future advances in LNP formulation design that allow a better balance between efficient mAb expression and minimal immunogenicity are likely critical for the clinical development of LNP-based mAb delivery strategies.

Overall, this study demonstrates that using mRNA/LNPs to deliver neutralising mAbs is a tractable pathway to protection against respiratory viruses such as SARS-CoV-2 and influenza. Serum concentrations of mAbs obtained by mRNA/LNP delivery were comparable to those obtainable with direct administration of recombinant proteins, although further work is needed to identify LNP formulations best suited for maximising the accumulation, retention and potentially mucosal localisation of antiviral mAbs. The rapid manufacturing and clinical development pathways for mRNA/LNP technologies make it particularly well-suited to strengthening pandemic preparedness, with efficient and cost-effective delivery of antiviral antibodies or biologics constituting a readily exploitable pathway to better protect human populations.

## MATERIALS AND METHODS

### Animal details and ethics statement

C57BL/6 and k18-hACE2 mice were bred at Australian Bio Resources (ABR) or Ozgene (WA, Australia) and housed at the Peter Doherty Institute’s Biological Resource Facility (BRF). All procedures involving animals and live SARS-CoV-2 were conducted in an OCTR-approved Physical Containment Level 3 (PC3) facility at the Centre for AgriBiosciences (AgriBio). All animal studies and related experimental procedures were approved by the University of Melbourne Animal Ethics Committee (No. 24909 and 22954). Female mice at 6 – 12 weeks old (n = 5 per group) were used for all studies.

### mAb mRNA design and synthesis

All mRNAs were synthesised at BASE mRNA facility (the University of Queensland), according to the method published previously.^22^ Briefly, a synthetic DNA encoding either heavy chain (HC), light chain (LC), or both heavy and light chains linked by P2A peptide (HC-P2A-LC) was obtained from Integrated DNA Technologies (Singapore). The DNA was then PCR amplified and used as templates for in vitro transcription of mRNA using T7 RNA polymerase (New England Biolab) with all uridine residues substituted by N1-methylpseudouridine (BOC Sciences). The mRNA products were purified using Monarch RNA Cleanup Kit (NEB), eluted in 1 mM sodium citrate and filtered sterile with 0.22-μm syringe filter.

### Preparation of mAb mRNA/LNPs

MAb mRNA/LNPs were prepared using the microfluidic mixing method.^16,20^ Briefly, lipids (DC Chemicals), including ionizable lipids (ALC-0315, Dlin-MC3-DMA), cationic lipid (DOTAP), helper lipid (DSPC), cholesterol, and PEG-lipids (ALC-0159, DMG-PEG2000) at specific molecular ratios were dissolved in ethanol where applicable. The mRNAs encoding HC and LC at 1:1 molar ratio admix or HC-P2A-LC were diluted in acetate buffer (pH 4.0). The two solutions were injected into a microfluidic Ignite Cartridge using a NanoAssemblr Ignite system (Precision NanoSystems) at an aqueous to ethanol ratio of 3:1 (vol./vol.), a total flow rate of 8 mL min^-1^, and a flow rate ratio of 3:1. The formulations were subsequently purified by dialysis against 10% sucrose in TBS buffer (Sigma Aldrich) over 18 – 20 h at room temperature. After that, the mRNA/LNPs were filtered using a Durapore 0.45-μm PVDF membrane (Merck) before being stored at –80 °C until use.

### Characterisation of mAb mRNA/LNPs

*Dynamic Light Scattering (DLS):* Dynamic diameter and zeta potential of mRNA LNPs (50 – 100 µg mL^-1^ in PBS) were measured by DLS using a Malvern Zetasizer Nano Series. *Encapsulation efficiency (EE):* the total amount of mRNA and unencapsulated mRNA in LNPs (with or without the addition of 1 µl of 10% Triton X-100 and continuous mixing at 300 rpm, 37 °C in 8 mins) were quantified using a Quant-iT™ RiboGreen^®^ RNA assay kit (Thermo Fisher Scientific) according to manufacturer guidelines. The EE of mRNA in LNPs was calculated with the following formula: EE% = (total mRNA – unencapsulated mRNA) ÷ total mRNA × 100.

### In vitro expression of mRNA-delivered mAbs

HEK293T cells were seeded in 6-well plates at ∼10^6^ cells per well and incubated overnight. The cells were then transfected with LNPs delivered HC and LC mRNA admix (1:1 molar ratio) or HC-P2A-LC mRNAs using Lipofectamine 3000 Transfection Reagents (Thermo Fisher Scientific) in Opti-MEM I Reduced Serum Medium (Thermo Fisher Scientific). Four hours later, the medium was replaced with DMEM (10% fetal calf serum (FCS), 1× penicillin-streptomycin-glutamine (PSG)). The supernatant was collected at 48h after transfection and the mAb expression was detected using ELISA as described below.

### In vivo expression and PK/PD studies of mAb mRNA/LNPs

C57BL/6 mice were i.v. administered with 200 μL LNPs delivered HC and LC admix or HC-P2A-LC mRNAs into the tail veins. At 48h post injection, blood samples were taken by submandibular or cardiac bleeds and serum was isolated by centrifugation. The expression of mRNA-delivered mAbs was confirmed by ELISA.

In PK studies, C57BL/6 mice were i.v. administered with 200 μL mAb mRNA/LNPs or mAb protein at 10, 5, or 1 μg in PBS. Time-series samples of mouse serum were collected. The mAb levels over time were quantified by ELISA, which were then used to estimate PK parameters for each dose event by using non-compartmental analysis.

In PD studies, two LNP formulations (ALC-0315 and MC3/DOTAP) delivering PDI204 HC-P2A-LC mRNA at a dose of 5 μg were administered into C57BL/6 mice via three different routes (i.v., i.m., and i.n.). At 48h post administration, serum, BALF and nasal wash samples were collected. For BALF and nasal wash collection, a small incision in the trachea was occupied to allow the insertion of a 20G cannula. The lungs were washed three times with 600 μL PBS via the cannula to harvest BALF samples. Nasal washes were taken by washing the nasal cavity three times with 200 μL PBS. The mAb expression in the samples was calculated using ELISA assays.

### ELISA for mAb detection

MAb expression in cell supernatant, mouse serum, BALF, and nasal washes was detected by a direct ELISA. The 96-well MaxiSorp plates were coated with 100 μL of antigens (either SARS-CoV-2 spike or A/California/04/2009 HA protein) at 2 μg mL^-1^ in PBS overnight at 4 °C. The plates were then blocked with 1% FCS for 1 h at room temperature. All serum samples were diluted in blocking solution at 1:10, while BALF samples and cell supernatant started as neat, followed by a serial fourfold dilution. Nasal washes were used as neat. The sample dilutions were added to the plates and incubated for 2 h at room temperature. The plates were next placed with anti-human secondary antibodies conjugated with HRP at 1:15,000 dilution in the blocking solution for 1 h at room temperature. The plates were then developed with 80 μL TMB in 8 mins and stopped with 50 μL of 0.16 M sulfuric acid. Absorbance was measured at 450 nm. mAb concentrations in the serum and BALF samples were quantified based on standard curves of mAb protein dilutions while ELISA endpoint titres were calculated as the reciprocal of sample dilution giving a signal 2× above background. For nasal washes, mAb levels were reported as optical density at 450 nm (OD450).

### ELISA to detect anti-mAb antibodies

Antibodies against mAbs in mouse serum at days 1, 7, and 14 following i.v. administration of mAb mRNA/LNPs and mAb proteins were calculated using a direct ELISA assay as described previously. Here, the plates were coated with mAb proteins at 2 μg mL^-1^ in PBS overnight at 4 °C. Serum samples were diluted in blocking solutions at 1:10, followed by a fourfold serial dilution. HRP-conjugated anti-mouse secondary antibody at 1:15,000 dilution was used.

### Microneutralisation assay

Neutralisation activity of PDI204 mAbs in mouse serum against the ancestral SARS-CoV-2 strain was performed as previously described.^23^ Briefly, SARS-CoV-2 isolate CoV/Australia/VIC31/2020 was passaged in Vero cells and stored at −80 °C.^24^ Infectivity of virus stocks was then determined by titration on HAT-24 cells (a clone of transduced HEK293T cells stably expressing human ACE2 and TMPRSS2).^25^ In a 96-well flat-bottom plate, virus stocks were serially diluted five-fold (1:5–1:78125) in DMEM with 1 μg mL^−1^ TPCK trypsin, added with 60000 freshly trypsinized HAT-24 cells per well and incubated at 37 °C. After 24 h, 10 μL of alamarBlue Cell Viability Reagent (ThermoFisher) was added into each well and incubated at 37 °C for 1 h. The reaction was then stopped with 1% SDS and read on a FLUOstar Omega plate reader (excitation wavelength 560 nm, emission wavelength 590 nm). The relative fluorescent units (RFU) measured were used to calculate % viability (“sample” ÷ “no virus control” × 100), which was then plotted as a sigmoidal dose–response curve on Graphpad Prism to obtain the virus dilution that induces 50% cell death (50% lethal infectious dose; LD50). Virus stocks were titrated in quintuplicate in three independent experiments to obtain mean LD50 values.

To determine serum neutralisation activity, heat-inactivated mouse serum samples were diluted 2.5-fold (1:50–1:30517) in duplicate and incubated with SARS-CoV-2 virus at a final concentration of 2 × LD50 at 37 °C for 1 h. Next, 60 000 freshly trypsinized HAT-24 cells in DMEM with 5% FCS were added and incubated at 37 °C. “Cells only” and “Virus+Cells” controls were included to represent 0% and 100% infectivity respectively. After 24 h, 10 μL of alamarBlue Cell Viability Reagent (ThermoFisher) was added into each well and incubated at 37 °C for 1 h. The reaction was then stopped with 1% SDS and read on a FLUOstar Omega plate reader. The relative fluorescent units (RFU) measured were used to calculate % neutralization with the following formula: (“Sample” – “Virus+Cells”) ÷ (“Cells only” – “Virus+Cells”) × 100. IC50 values were determined using four-parameter non-linear regression in GraphPad Prism with curve fit constrained to have a minimum of 0% and a maximum of 100% neutralisation.

### Prophylaxis studies of mAb mRNA/LNPs against SARS-CoV-2 infection in mice

Prophylactic efficacy of PDI204 mRNA/LNPs versus proteins was assessed in K18-hACE2 mice. Briefly, K18-hACE2 mice were i.v. injected with 200 μL of PDI204 mRNA/LNPs at 10, 5, or 1 μg, protein or irrelevant mAb mRNA/LNPs at 10 μg. Seven days post treatment, mice were lightly anesthetized (isoflurane) and intranasally infected with 10^4^ 50% Tissue Culture Infectious Dose (TCID50) of SARS-CoV-2 Omicron BA.1 (strain SARS-CoV-2/Australia/NSW/RPAH-1933/2021) in 50 μl PBS. On day 4 post-infection, mice were humanely killed. Nasal turbinates and lungs were collected and homogenised using an Omni tissue homogenizer in either 1 mL or 2 mL, respectively of PBS containing 50 U/mL of Penicillin, 50 μg/mL Streptomycin and 4 µg/mL Amphotericin B (Gibco). Homogenates were clarified by centrifugation at 2,000 x g for 10 minutes before virus quantification by a TCID50 assay. Briefly, clarified tissue homogenates were serially diluted (1:10) on confluent monolayers on VeroE6 cells over-expressing TMPRSS2 in DMEM containing 50 U/mL of Penicillin, 50 μg/mL Streptomycin, 2 mM Glutamax (Gibco) and 1 µg/mL trypsin (Worthington Biochemical) in quadruplicate. Plates were incubated at 37 °C supplied with 5% CO_2_ for 4 days before measuring cytopathic effect under a light microscope. TCID_50_ was determined using the Reed-Muench method. Virus titers are expressed as mean log_10_ TCID_50_/mL.

### Prophylaxis studies of mAb mRNA/LNPs against influenza infection in mice

Prophylactic efficacy of HV-B10 mRNA/LNPs versus proteins was assessed in murine infection models as described previously.^14,20^ C57BL/6 mice were i.v. injected with 200 μL of HV-B10 mRNA/LNPs or protein at 10, 5, or 1 μg, or irrelevant mAb mRNA/LNPs control (10 μg) at 24 h prior to viral infection. The mice were intranasally infected with 50 μL of A/California/04/2009 virus at a lethal dose of 100 pfu in PBS. The mice were monitored for clinical signs and weight loss for 14 days post-infection. Mice were euthanised if 20% of their initial weight was lost.

### Statistical analysis

Data were presented as median ± interquartile range (IQR) (n = 5). Statistical analyses were performed using GraphPad Prism version 10. For all analyses comparing multiple groups, a Kruskal-Wallis test followed by post-hoc Dunn’s multiple comparisons test was used. *p* values of less than 0.05 (*p* < 0.05) were considered to be significant for all statistical tests.

## ACKNOWLEDGMENTS

Some portions of the figures were created with BioRender (BioRender.com). All mAb mRNA constructs used in this study were synthesised by BASE mRNA Facility at the University of Queensland. The authors acknowledge the facilities and technical assistance of the Bioresources Facility (BRF) at the University of Melbourne and the Centre for AgriBiosciences (AgriBio) at La Trobe University. This work was supported by Australian Medical Research Future Fund grants 2005544 and 2013870, Australian National Health and Medical Research Council Investigator grants (W.S.L, H.-X.T., and A.K.W.), and Therapeutic Innovation Australia Pipeline Accelerator Voucher Program (M.N.V.).

## SUPPORTING INFORMATION

**Table S1:**
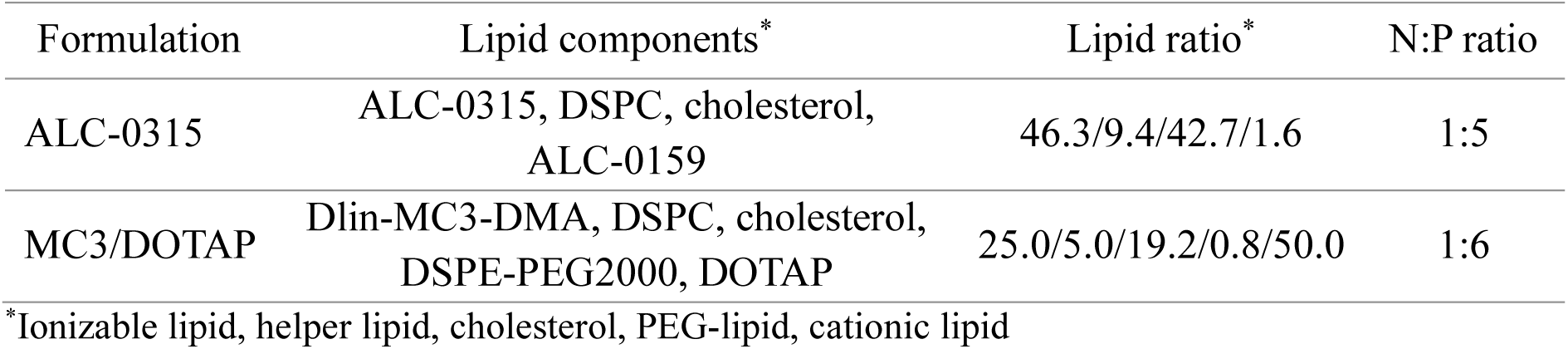
Formulation of mAb mRNA/LNPs.

**Table S2:** Characterisation of mRNA/LNPs.

| Formulation | mRNA | Diameter (nm) | PdI | Zeta potential (mV) | Encapsulation efficiency (%) |
| --- | --- | --- | --- | --- | --- |
| ALC-0315 | HC/LC admix | 62.3 ± 1.5 | 0.102 | - 8.31 | 89.3 |
|  | HC-P2A-LC | 64.1 ± 1.3 | 0.116 | - 8.09 | 90.6 |
| MC3-DOTAP | HC-P2A-LC | 121.7 ± 4.2 | 0.146 | + 15.2 | 96.5 |

